# HitAnno: Atlas-level cell type annotation based on scATAC-seq data via a hierarchical language model

**DOI:** 10.64898/2026.03.10.710729

**Authors:** Zian Wang, Xiaoyang Chen, Xuejian Cui, Zijing Gao, Zhen Li, Keyi Li, Rui Jiang

**Author notes:** These authors contributed equally: Zian Wang, Xiaoyang Chen.

## Abstract

The single-cell assay for transposase-accessible chromatin using sequencing (scATAC-seq) has emerged as a core technology for dissecting cellular epigenomic heterogeneity and gene regulatory programs. With the emergence of atlas-level scATAC-seq datasets, cell type annotation increasingly faces challenges arising from unprecedented data scale and increased cell-type diversity, which together place stringent demands on model reliability and robustness. Here, we introduce HitAnno, a hierarchical language model capable of accurate and scalable cell type annotation in atlas-level scATAC-seq data. Leveraging selected cell-type-specific peaks to construct “cell sentences”, HitAnno employs a two-level attention mechanism that captures accessibility profiles hierarchically. Extensive evaluations show that HitAnno robustly annotates both major and rare cell types across multiple settings, including intra-dataset, cross-donor and inter-dataset annotation. The hierarchical attention mechanisms of the model reveal co-accessibility patterns among peaks and dependencies across higher-order peak sets, ensuring an interpretable annotation process. Training on a 31-cell-type human atlas, HitAnno can directly annotate new query datasets without retraining and is accessible through an online interface. Our model identifies heterogeneous subgroups within mixed labeled cells from unseen datasets, demonstrating its potential to assist researchers in refining existing cell atlases.

## Introduction

The advent of the single-cell assay for transposase-accessible chromatin using sequencing (scATAC-seq) opens new avenues to investigate the epigenomic regulatory landscape at single-cell resolution^1-3^, providing insight into delineating cell differentiation^4^, characterizing developmental dynamics^5^, and identifying disease-associated variation^6^. As a prerequisite for these downstream analysis tasks, the accurate annotation of cell types based on scATAC-seq data reveals cellular heterogeneity and enables the analysis of cell-type-specific accessibility profiles, thereby facilitating our understanding of regulatory mechanisms across different tissues. In practice, the annotation procedure typically relies on unsupervised clustering based on accessibility profiles, followed by manual labeling using cell markers specific to clusters^7-9^. This procedure is often tedious, labor-intensive and difficult to reproduce, as manual labeling depends on non-standardized cell labels and expert knowledge^10^, which highlights the urgent need for automated cell type annotation methods.

In recent years, the increasing availability of high-quality, well-annotated scATAC-seq datasets drives the development of supervised methods for automatic cell type annotation^11, 12^, leading to computational methods that adopt diverse modeling strategies. Several representative methods include: (1) probabilistic-based methods, such as EpiAnno, which employs a Bayesian neural network to project cells into a latent space governed by a Gaussian mixture distribution, and allows for annotation based on learned probabilistic assignments^13^; (2) gene-activity-based methods, such as Cellcano, which uses gene activity scores derived from chromatin accessibility profiles and employs a two-round training strategy^14^; (3) sequence-based methods, such as SANGO, which incorporates modeling of DNA sequences surrounding accessible regions and leverages a two-stage deep learning approach^15^. Collectively, these methods demonstrate the feasibility of supervised cell type annotation on scATAC-seq data.

However, as scATAC-seq datasets continue to grow in size and diversity, cell type annotation increasingly requires extension to large-scale atlases, which are characterized by complex and heterogeneous cell populations, and existing methods still face several challenges. First, the massive cell numbers of atlas-level datasets, combined with the intrinsic characteristics of scATAC-seq data, including high dimensionality, extreme sparsity, and a near-binary signal structure, together place stringent demands on model scalability and robustness. Second, the expansion of cell type diversity and imbalance in cell type abundance introduces substantial bias during supervised learning, as learned representations tend to be dominated by major cell types, hindering the discrimination of rare populations. Third, interpretability on the annotation process is essential to ensure that models capture biologically meaningful patterns rather than dataset-specific noise, which is critical for the reliable application on atlas-level datasets.

To address these challenges, we developed HitAnno, a hierarchical language model based on transformer architecture capable of atlas-level cell type annotation on scATAC-seq data. HitAnno represents each cell as a structured “cell sentence”, constructed from accessibility profiles on specific peaks with consideration of both major and rare cell types. Based on cell sentences, HitAnno employs a two-level attention mechanism that integrates accessibility profiles from individual peaks to higher-order peak sets, enabling scalable extraction of cell-type-specific patterns while explicitly capturing peak interactions at hierarchical levels. In intra-dataset evaluations, HitAnno consistently achieved accurate annotation of both major and rare cell types, demonstrating robust performance under challenging settings. The hierarchical attention mechanism further enabled interpretable analysis by capturing both local co-accessibility patterns among peaks and global dependencies across peak sets. In more practical annotation tasks, HitAnno maintained reliable performance in cross-donor and inter-dataset annotations, demonstrating robustness across heterogeneous data sources. Finally, scaling HitAnno to atlas-level training allowed direct annotation of new query datasets without retraining, supporting its deployment as a reusable annotation model. We further packaged this atlas-level model into an online annotation platform and demonstrated its ability to separate biologically coherent subgroups within mixed-labeled populations in new query datasets.

## Results

### Overview of HitAnno

HitAnno is a supervised method for accurate and scalable cell type annotation based on scATAC-seq data. Our method is designed from the perspective of “chromatin accessibility landscape as a language”, in which a cell is described as a long sentence covering candidate peaks, along with their accessibility states. To make this intuitive formulation practical, we introduce a hierarchical language model, which divides a long cell sentence into several clauses (analogous to functional sub-sentences), each corresponding to a cell type and using only a specific subset of peaks. As illustrated in Fig. 1, this model consists of three modules: tokenization, representation, and annotation.

**Fig. 1.**
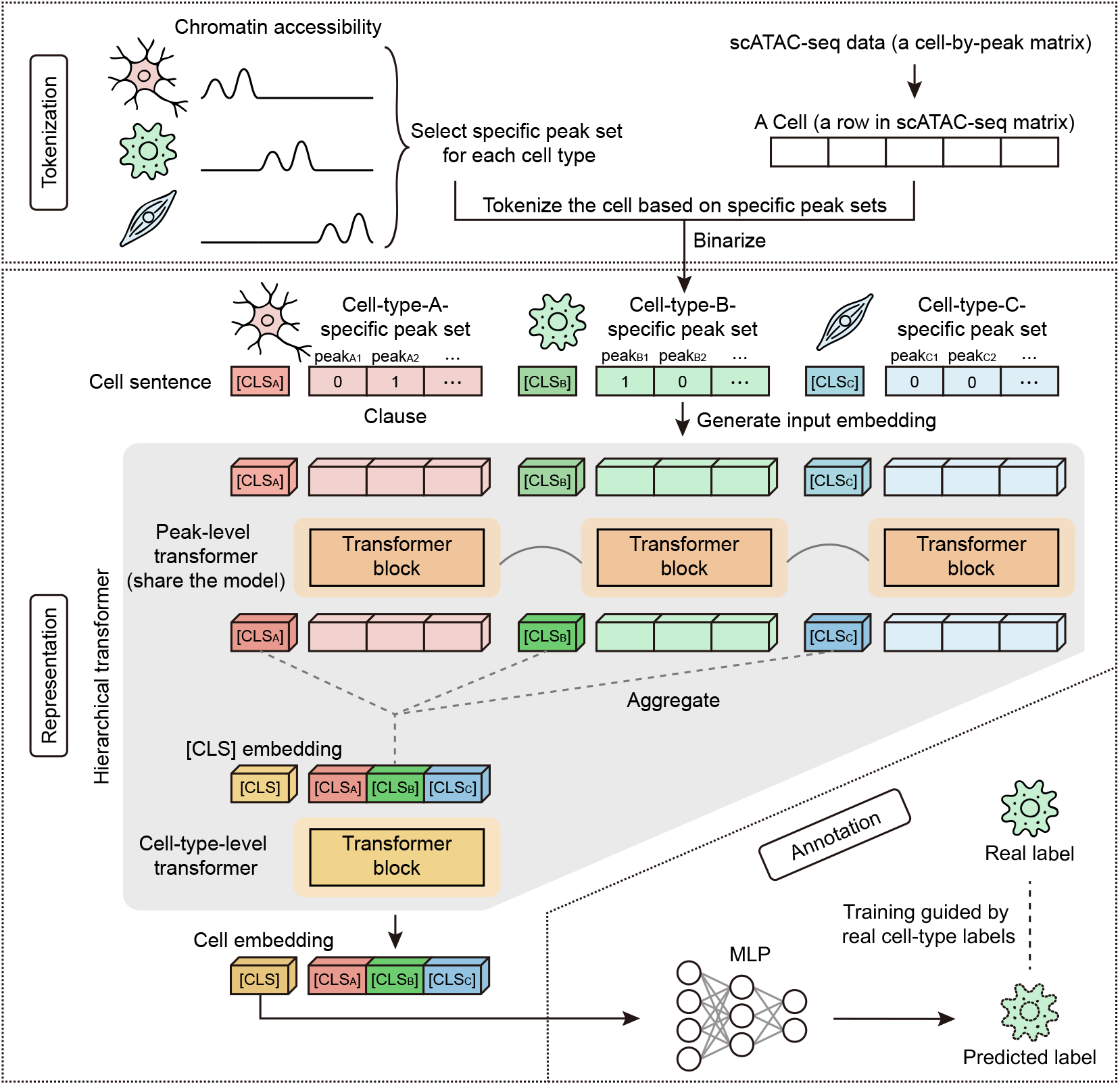
Overview of HitAnno. This method comprises three modules: tokenization, representation, and annotation. The tokenization module identifies specific peak set for each cell type and tokenizes the cell into a cell sentence. In the representation module, the cell sentence is embedded by combining binary accessibility values with peak indices, producing an input embedding. Then, a hierarchical transformer employs a two-level attention mechanism to extract a cell embedding. Finally, the annotation module utilizes an MLP-based classifier to generate predicted cell type probabilities from the cell embedding.

The tokenization module converts the chromatin accessibility landscape of a cell into a cell sentence composed of several clauses, as follows. First, a peak selection step identifies for each cell type a set of specific peaks, forming a specialized vocabulary. Then, a cell tokenization step describes each cell as a sequence of clauses. A clause corresponds to a cell type, either major or rare, and is represented as a sequence of peaks specific to the cell type, prepending a class token [CLS_CT_] that marks the clause (Methods). All clauses have equal length to prevent the model from focusing primarily on major cell types.

The representation module captures interactions between peaks in a cell sentence and produces an embedding for the corresponding cell. Given the large number of peaks in all clauses, it is impractical to capture interactions between peaks using a vanilla transformer. We therefore designed a hierarchical transformer architecture by adopting the idea of divide-and-conquer, as follows. First, cell sentence is converted to input embedding. Particularly, the index and binary accessibility values of each peak are represented by separate embeddings and then added to produce the input embedding. Next, a peak-level transformer is applied to each clause to capture interactions between peaks within each cell type and produces embeddings for the [CLS_CT_] tokens. Then, a cell-type-level transformer is applied to capture dependencies between cell types and produces the embedding for the [CLS] tokens. This embedding is used to represent the corresponding cell. Details of this module are provided in Methods.

The annotation module accepts a cell embedding as input and resorts to a multi-layer perceptron (MLP) to calculate a probability distribution of predicted cell types as output.

### HitAnno achieves accurate and robust intra-dataset annotation

We first evaluated HitAnno in a relatively ideal intra-dataset setting, where reference cells and query cells are drawn from the same dataset. To achieve this objective, we collected 12 scATAC-seq datasets spanning diverse sequencing depth, platform, and cell population heterogeneity (Supplementary Table 1). For each dataset, we performed five-fold cross-validation, evaluating the performance of HitAnno using three metrics, accuracy, macro F1 score (macro-F1), and Cohen’s kappa score (*κ*). HitAnno was compared with state-of-the-art scATAC-seq annotation methods (SANGO^15^, Cellcano^14^, and EpiAnno^13^), and previously benchmarked general-purpose classifiers (MLP^16^ and Principal Component Analysis and Support Vector Machine, PCA+SVM^10^). Details of the evaluation criteria and baseline methods are given in Methods.

As shown in Fig. 2a, our method consistently outperforms the five baseline methods across all twelve datasets, achieving an average accuracy of 89.74%, which is 6.91% higher than the second-best method, SANGO. In addition, our model achieves improvements of 8.11% and 9.25% over SANGO in macro-F1 and *κ*, respectively. On datasets where baseline methods already achieve high accuracy (e.g., Domcke2020_thymus, thymus dataset from Domcke et al., 2020^17^; Zhang2021_liver, liver dataset from Zhang et al., 2021^18^), HitAnno still demonstrates clear advantages on metrics that strongly reflect performance across heterogeneous cell populations. For example, on Domcke2020_thymus, HitAnno improves macro-F1 and *κ* by 7.16% and 6.84%, respectively, indicating superior discrimination of diverse cell populations, particularly rare cell types. On more challenging datasets, characterized by high cell type diversity (e.g., Zhang2021_artery and Zhang2021_lung), HitAnno exhibits even more pronounced gains, with macro-F1 gains of up to 13.94%. Together, these consistent improvements across datasets of varying difficulty demonstrate the effectiveness of our model design. By constructing cell sentences that incorporate cell-type-specific peaks, HitAnno is able to robustly capture discriminative chromatin accessibility patterns for both major and rare cell populations.

**Fig. 2.**
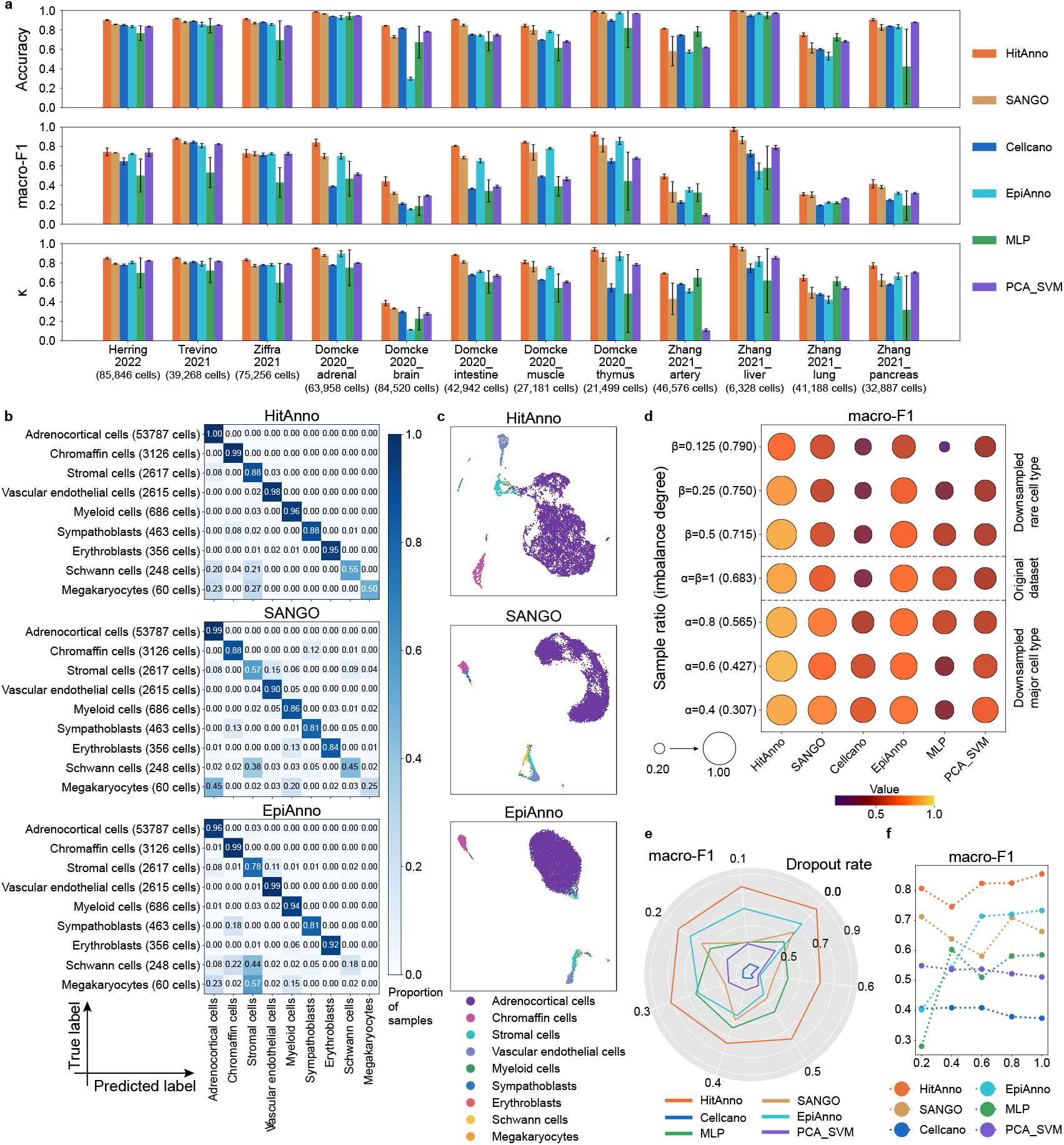
HitAnno achieves accurate and consistent intra-dataset annotation. **a**, Accuracy, macro-F1, and *κ* across five-fold cross-validation on 12 datasets, comparing HitAnno with five baseline methods. Bar heights indicate average performance across folds, and error bars represent 95% confidence intervals. **b**, Confusion matrices of the top three performing methods (ranked by macro-F1) on Domcke2020_adrenal dataset, showing per-cell-type annotation performance, with the number of cells indicated in parentheses. **c**, UMAP visualization of cell embeddings learned by the top three methods on Domcke2020_adrenal dataset, colored by real cell type labels. **d-f**, Macro-F1 of HitAnno and five baseline methods under varying levels of cell type imbalance (**d**; here, *α* and *β* denote the retention fractions of cells for major and rare cell types, respectively), dropout rates (**e**), and downsampling ratios (**f**).

We further examined the confusion matrix of Domcke2020_adrenal (Fig. 2b) to provide a more fine-grained view of how the performance gains of HitAnno are distributed across cell types. For the major cell type, adrenocortical cells, HitAnno achieves near-perfect annotation. As cell type abundance decreases, HitAnno remains robust across rare populations, with six out of eight rare cell types achieving F1 scores above 0.88. Notably, even for the two rarest cell types, HitAnno preserves relatively stable performance, whereas baseline methods often exhibit pronounced declines. Consistent with these observations, UMAP visualization of the top three performing methods on this dataset (Fig. 2c) further demonstrates the superior performance of HitAnno. The embedding space learned by HitAnno forms tight, well-separated clusters for different cell types, reflecting the ability of the model to capture discriminative cell-type-specific patterns.

Accurate cell type annotation on scATAC-seq data requires robust models, which in real-world applications often present challenging conditions, such as differences in cell type proportions, extremely sparse accessibility, and reduced data availability. Accordingly, we conducted three types of experiments: varying degrees of cell type imbalance^19, 20^ (Supplementary Note 1), simulated dropout, and downsampling of cell numbers. HitAnno shows consistent robustness advantages over all five baselines across these conditions. Under extreme cell type imbalance (0.79), our model achieves a 12.5% improvement in macro-F1 over the second-best method (Fig. 2d; Supplementary Figs. 1). Similar advantages are observed under dropout conditions, where HitAnno attains an average macro-F1 that is 15.33% higher than that of the second-best method (Fig. 2e; Supplementary Figs. 2). Across downsampling scenarios, HitAnno not only maintains superior accuracy, macro-F1, and *κ*, but also shows remarkable stability, whereas most baselines are unstable and consistently underperform (Fig. 2f; Supplementary Figs. 3).

**Fig. 3.**
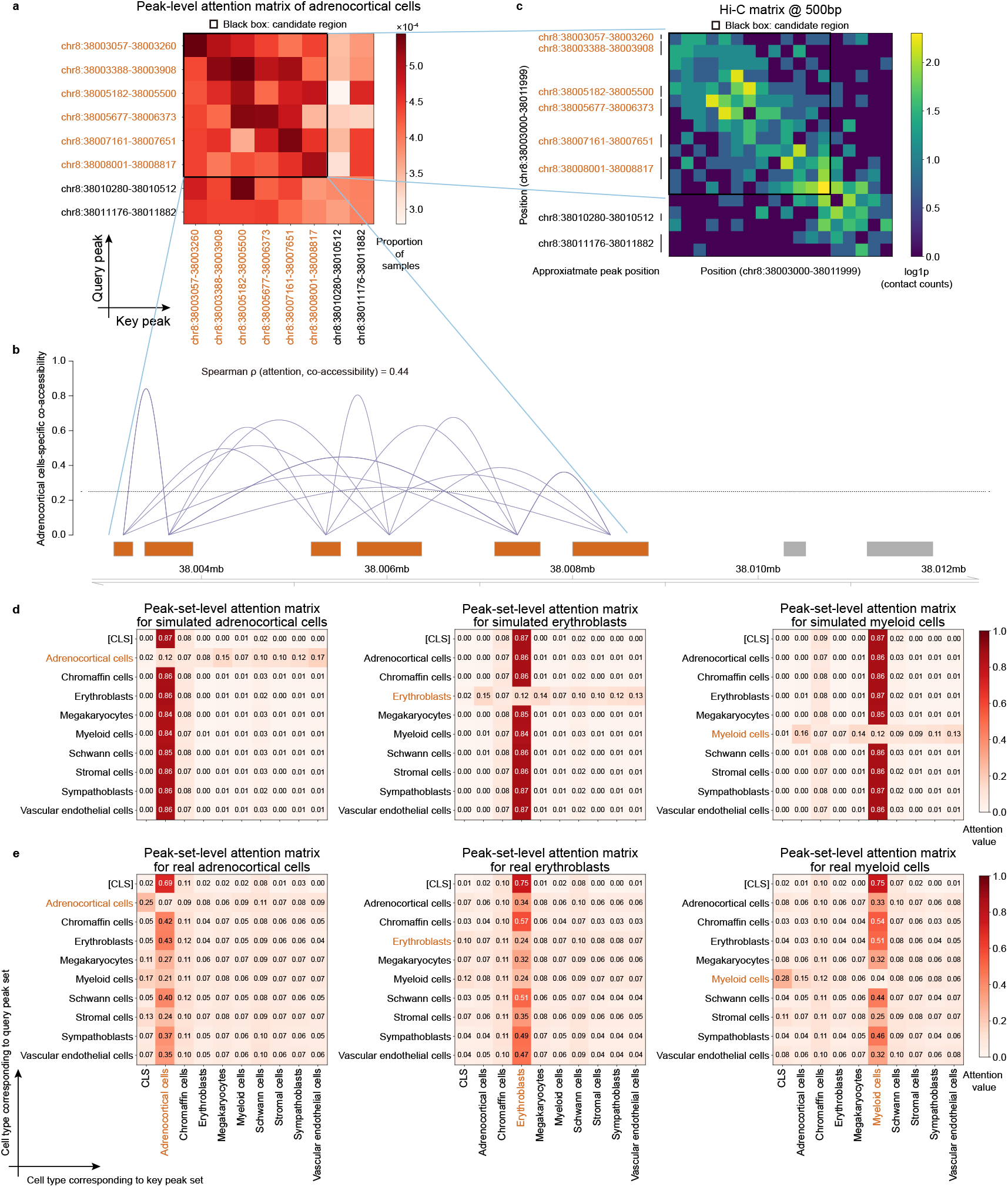
HitAnno captures interpretable patterns through hierarchical attention. **a**, A localized region of the peak-level attention matrix for adrenocortical cells in the Domcke2020_adrenal dataset. **b**, Co-accessibility map predicted by Cicero for adrenocortical cells, with a co-accessibility score threshold of 0.25. **c**, Hi-C contact map for the same genomic locus from human adrenal tissue. The candidate region identified by HitAnno is marked in black. **d-e**, Peak-set-level attention matrices computed using simulated inputs (**d**) and real dataset (**e**).

To assess the contribution of key architectural components, we conducted ablation studies by simplifying the hierarchical transformer or replacing cell-type-specific peak selection with global peak selection. As detailed in Supplementary Fig. 4, both modifications led to consistent performance degradation across all metrics, confirming the importance of these design choices.

**Fig. 4.**
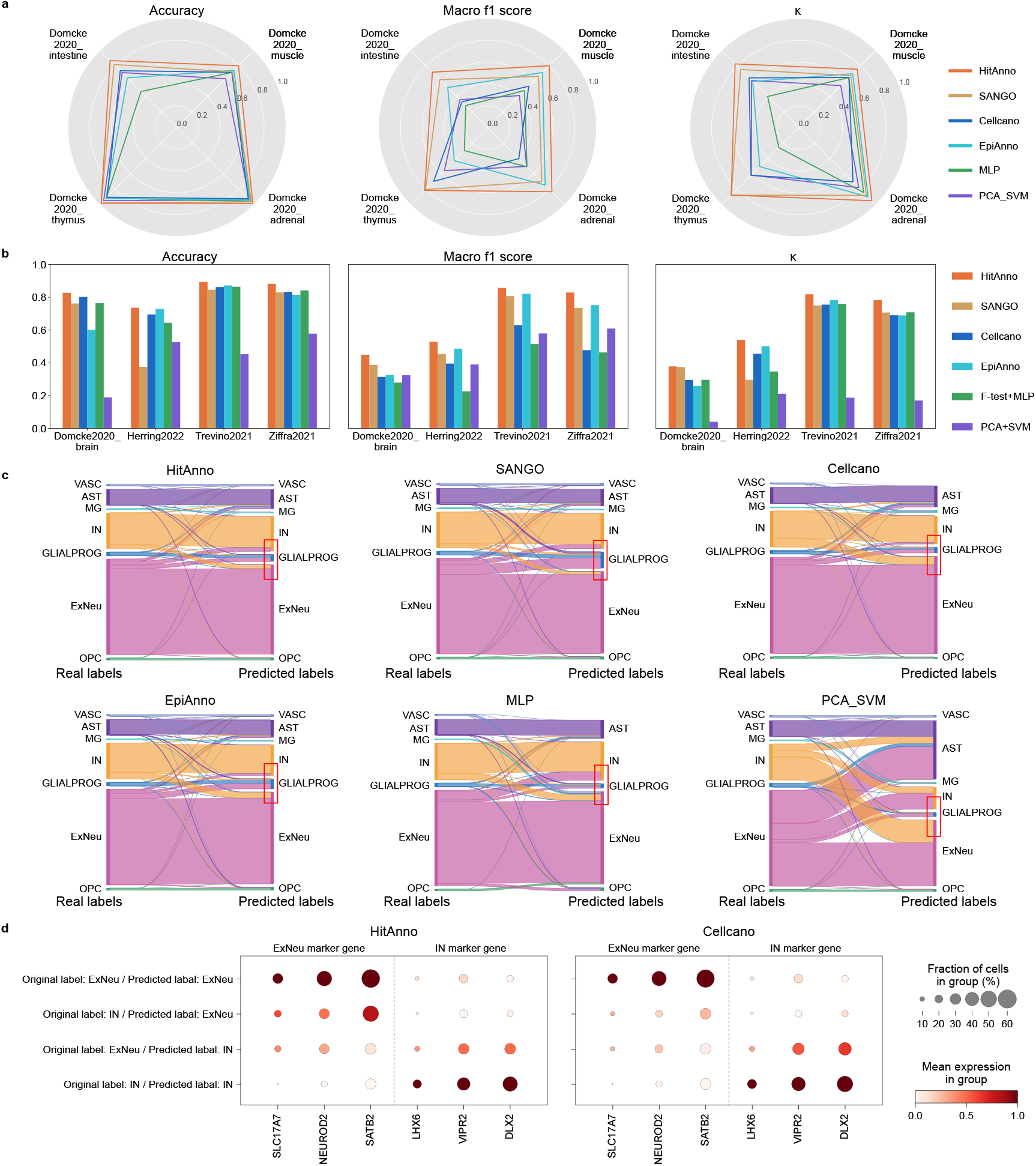
HitAnno demonstrates generalization across donors and datasets. **a**, Accuracy, macro-F1, and *κ* for cross-donor annotation on four datasets. **b**, Performance comparison of HitAnno and five baseline methods in inter-dataset annotation tasks, evaluated on four independent query datasets using accuracy, macro-F1, and *κ*. **c**, Sankey plots illustrating cell type annotation flows on the Trevino2021 dataset for HitAnno and five baseline methods. Real cell type labels are shown on the left, and predicted labels on the right. **d**, Gene activity scores of marker genes for ExNeu and IN in four categories of cells based on original and predicted labels, shown for the top two performing methods in the inter-dataset annotation task. Dot color represents the average gene activity score, and dot size reflects the fraction of cells in each category with non-zero activity for the corresponding marker gene.

### HitAnno captures interpretable patterns through hierarchical attention

In addition to the strong intra-dataset performance of HitAnno, we further investigated the interpretability of the trained model. Enabled by its hierarchical transformer architecture, HitAnno supports interpretation based on attention mechanisms at both peak and peak-set levels. We therefore examined attention patterns across different hierarchical levels to characterize their roles in cell type annotation.

At the peak level, HitAnno attention reveals interactions among peaks within local regions, reflecting local co-accessibility patterns relevant to cell identity. As an illustrative example (Fig. 3a), one high-attention region spans the STAR locus in adrenocortical cells from the Domcke2020_adrenal dataset, a key regulator of steroidogenesis and a well-established marker of adrenocortical cell identity^21, 22^. We also examined results on the same region using Cicero^23^, which identifies co-accessible pairs of DNA elements using single-cell chromatin accessibility data. As shown in Fig. 3b, multiple peaks within this region exhibited clear co-accessibility. Moreover, peak pairs with detectable co-accessibility tended to display higher attention scores, indicating a moderate positive association between attention patterns learned by HitAnno and co-accessibility signals inferred by an independent, data-driven method.

Moreover, we examined the same genomic locus from a complementary perspective using three-dimensional genome organization data. Comparison with Hi-C data from human adrenal tissue shows that the local contact pattern at this locus qualitatively resembles the high-attention region in the peak-level attention matrix (Fig. 3c). Elevated contact frequencies are observed near the peak positions, further supporting the existence of functional interactions. Together, evidence from both Cicero co-accessibility and Hi-C contacts demonstrates that HitAnno attention captures biologically meaningful relationships among peaks, providing peak-level interpretability.

At peak-set level, HitAnno captures higher-order dependencies, with each cell type exhibiting a distinct and interpretable attention pattern. To illustrate the ideal attention patterns of HitAnno, we first examined peak-set-level attention matrices under controlled simulated inputs, providing a reference for interpreting the model in real scATAC-seq data. Specifically, we constructed simulated inputs in which only the peaks specific to a single cell type are set as accessible, while all other peaks are considered inaccessible. Under this setting, the model attention assigns higher attention weights to the accessible peak set which dominates the [CLS] token representation, indicating that the model assigns attention in a cell-type-discriminative manner at the peak-set level (Fig. 3d, Supplementary Fig. 5).

**Fig. 5.**
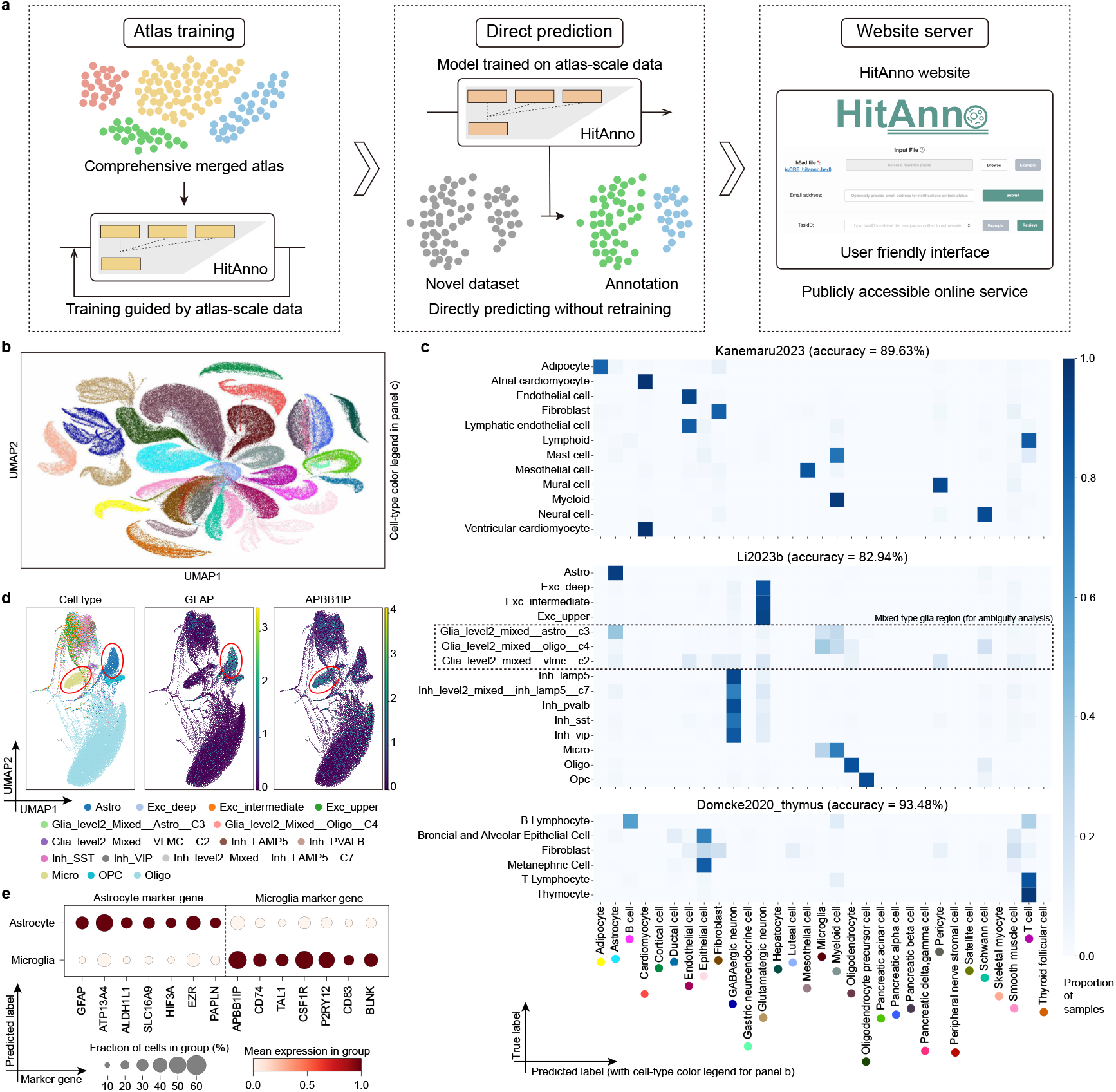
HitAnno supports scalable atlas-level training and annotation. **a**, Overview of the atlas-level training workflow. An integrated large-scale scATAC-seq atlas was used to train a unified HitAnno model. The trained model supports direct annotation of external datasets and has been deployed as an online tool. **b**, UMAP visualizations of the integrated large-scale scATAC-seq atlas, with all cells colored by their real cell type labels. **c**, Confusion matrices of HitAnno on three query datasets. The x-axis shows predicted cell type, and the y-axis shows real cell type. A black dashed box highlights the region corresponding to mixed-type glial populations, which is selected for downstream ambiguity analysis. **d**, UMAP visualizations of the Li2023b dataset. The left panel shows the cells colored by their real labels. The middle and right panels show the gene activity scores of the astrocyte and microglia marker genes, respectively. **e**, Gene activity scores of marker genes for astrocyte and microglia in two categories of cells based on predicted labels. Dot color represents the average gene activity score, and dot size reflects the fraction of cells in each category with non-zero activity for the corresponding marker gene.

More importantly, similar attention profiles are observed in real scATAC-seq data (Fig. 3e, Supplementary Fig. 6). In particular, attention weights are consistently concentrated on the peak sets corresponding to real cell type labels, consistent with the behavior observed under simulated inputs. This consistency further supports that HitAnno reliably focuses on patterns that are discriminative of cell identity across both simulated inputs and real data, forming stable global dependencies among peak sets.

### HitAnno shows reliable performance across donors and datasets

After evaluating intra-dataset annotation performance, we next assessed HitAnno in more practical annotation tasks, which reflects the real-world application of annotating cells from unseen donors or datasets using existing reference data. To this end, we first conducted cross-donor experiments on four datasets from Domcke2020, which exhibits batch effects between donors (Supplementary Fig. 7). Cells are split by donor, with each in turn used as the query and the remaining as the reference. Across four datasets, HitAnno consistently outperforms the five baseline methods (Fig. 4a). The largest improvement is 9.51% observed in macro-F1, indicating that HitAnno maintains robust annotation performance across diverse cell types despite batch effects. This robustness is consistent with the design of HitAnno, which captures cell-type-specific accessibility patterns and performs cell type prediction globally based on these patterns. The hierarchical separation of local feature extraction and global decision-making enables the model to consistently identify cell types across heterogeneous populations, thereby effectively mitigating the influence of batch effects.

We further evaluated the performance of HitAnno in practical inter-dataset annotation tasks on four large-scale brain scATAC-seq datasets: Domcke2020_brain, Herring2022, Trevino2021, and Ziffra2021. Each dataset in turn served as the query while the remaining three were used as references. These datasets span various human developmental periods. Trevino2021 comprises fetal cortical samples collected during mid-gestation^24^, while Herring2022 profiles prefrontal cortex development from fetal to adult stages^25^. Different sequencing technologies were employed across datasets: Domcke2020 used sci-ATAC-seq3^17^, and the other three adopted 10X ATAC-seq^24-26^. Such heterogeneity (Supplementary Fig. 8) makes inter-dataset annotation a useful way to assess the ability of the model to be applied to new, unseen datasets.

Across all four datasets and three evaluation metrics, HitAnno consistently achieves the best performance in the inter-dataset annotation task (Fig. 4b). On average, HitAnno achieves the strongest gains on macro-F1 (7.25%) and *κ* (7.20%). In contrast, the baseline methods show greater variability in performance, and no single method consistently ranks second across all metrics. Cellcano performs second best in accuracy, SANGO in macro-F1, and EpiAnno in *κ*. This variability reflects the lack of overall stability in existing approaches, whereas HitAnno remains robust across all metrics. Fig. 4c further compares cell type annotation flows on Trevino2021 using Sankey plots. HitAnno achieves notably higher accuracy for cell types such as IN (GABAergic neuron), GLIALPROG (glial progenitor cell), and ExNeu (excitatory neuron), with fewer annotation errors compared with other methods. This result further demonstrates the improved cell-type separability achieved by HitAnno across heterogeneous brain datasets.

The Trevino2021 dataset was further analyzed as the query dataset to investigate patterns of confusion between two major cell types, ExNeu and IN. We divided cells into four groups based on real and predicted labels. For each group, gene activity scores were computed from the original scATAC-seq data. We then assessed the gene activity patterns of these groups using ExNeu and IN marker genes curated from the literature^27, 28^. The results show that IN cells classified as ExNeu by HitAnno still exhibit high activity on ExNeu markers, whereas ExNeu cells classified as IN display gene activity patterns more similar to their original cell identity (Fig. 4d). In contrast, most baseline methods fail to retain clear marker signals in misclassified cells, or show substantially weaker patterns compared to HitAnno (Fig. 4d, Supplementary Fig. 9). During early neurodevelopment, chromatin accessibility patterns are progressively remodeled, creating challenges for distinguishing different cell types including ExNeu and IN cells^26^. Notably, HitAnno outperforms other methods in separating these cells, with predicted labels consistent with marker-based gene activity scores, further demonstrating its potential for practical applications in cross-dataset cell type annotation.

### HitAnno supports scalable atlas-level training and annotation

Motivated by its reliability across datasets, we extended HitAnno to an atlas-level setting (Fig. 5a), a scale at which most existing methods are not designed for or difficult to deploy due to scalability constraints. At atlas scale, cell type annotation requires models to handle both vastly increased cell numbers and substantially expanded cell-type diversity, placing demands on scalability and reliability. HitAnno is naturally suited to this setting due to its hierarchical language modeling formulation. For atlas-level datasets, HitAnno first selects cell-type-specific peaks to construct cell sentences, ensuring that the model captures specific accessibility patterns for each cell type. Each long cell sentence is then decomposed into multiple clauses, which are processed independently in parallel at peak level, enabling the model to scale effectively to atlases with large numbers of cells and diverse cell types.

We constructed a unified atlas-scale reference dataset by integrating adult brain and fetal scATAC-seq atlas (Li2023^29^ and Zhang2021), enabling robust cell-type annotation across a broader biological spectrum. After harmonizing peaks and cell-type labels, we constructed a unified reference atlas spanning 31 common cell types (Fig. 5b). The resulting model, trained on this large dataset, serves as a classifier capable of supporting new annotation tasks without the need for retraining.

We applied this trained atlas-level model directly to three external datasets, Kanemaru2023^30^, Li2023b^31^, and Domcke2020_thymus, to assess its performance. As shown in Fig. 5c, despite the 31-cell-type annotation challenge, HitAnno produces annotations that capture the majority of cell types in all external datasets. The overall annotation accuracies are 89.63%, 82.94%, and 93.48% for the three datasets, respectively. The model also captures hierarchical relationships among closely related subtypes; for example, atrial and ventricular cardiomyocytes are annotated under the broader cardiomyocyte category in Kanemaru2023, and Exc_deep, Exc_intermediate, and Exc_upper cells were grouped together as glutamatergic neurons in Li2023b. Together, these results demonstrate that the atlas-level HitAnno model can generalize across multiple external datasets, supporting its use as a scalable model for annotating large-scale scATAC-seq data.

We conducted a focused analysis on the Li2023b dataset to further evaluate the generalization of HitAnno. We observed that several glial cell types in this dataset were labeled as “mixed” in their cell-type annotations. These mixed glial populations are reported to have low-confidence labels in the original study, and the authors excluded them from subsequent analyses^31^. To assess the ability of HitAnno to distinguish cell types within the mixed glial populations, we compiled a set of known astrocyte and microglia markers from the literature and evaluated their gene activity scores in this dataset^27, 28^. We retained only markers that showed specific gene activity in confidently annotated astrocytes or microglia within this dataset to ensure marker specificity (Fig. 5d). We then examined the gene activity scores of these markers in cells predicted by HitAnno as astrocytes or microglia within the mixed glial populations. The results show that cells predicted as astrocytes exhibit higher gene activity scores of astrocyte-specific markers, whereas cells predicted as microglia show higher gene activity scores of microglia-specific markers (Fig. 5e, Supplementary Fig. 10, Supplementary Fig. 11). These observations indicate that the atlas-level HitAnno model can distinguish biologically meaningful subpopulations within challenging mixed glial populations, suggesting its potential utility for assisting researchers in resolving ambiguous cell types.

Finally, we deployed the atlas-level HitAnno model as an online tool to provide automated cell-type annotation (URL: https://health.tsinghua.edu.cn/hitanno). The web interface is user-friendly and includes detailed instructions and tutorials. Users can upload scATAC-seq data files to obtain direct access to the model online (Fig. 5a).

## Discussion

In this study, we present HitAnno, a hierarchical language model for accurate cell type annotation based on atlas-level scATAC-seq data. Through extensive evaluations, HitAnno demonstrates stable and consistent performance across intra-dataset annotations, reflecting a level of reliability that is not uniformly achieved by existing approaches. We further show that this capability extends to more practical and challenging scenarios, including cross-donor and inter-dataset annotation tasks, highlighting the utility of HitAnno in the presence of batch effects and cellular heterogeneity. Importantly, models trained at atlas scale can be directly deployed to annotate new query datasets without retraining, enabling the real-world application of HitAnno through an online interface.

By explicitly organizing peaks into cell-type-specific clauses, HitAnno constrains representation learning to focus on biologically meaningful accessibility patterns, rather than relying on the model to disentangle such structure from high-dimensional, heterogeneous inputs. This design not only improves robustness under batch effects and dataset shifts, but also enables efficient scaling to atlas-level datasets with large cell-type diversity. More broadly, this hierarchical formulation provides a general strategy for structuring representation learning in large-scale heterogeneous sequencing datasets beyond scATAC-seq.

Despite the promising potential of HitAnno across diverse tasks, several areas warrant further exploration and refinement. First, the model relies on predefined peak sets, which may limit discovery of novel regulatory regions. Moreover, with the increasing availability of multi-omics data^32-34^, HitAnno could be extended to incorporate additional modalities such as scRNA-seq and scHi-C, facilitating a more comprehensive modeling of cellular states^35, 36^. Finally, as pretrained generative models become increasingly prominent^37-40^, cell type annotation is evolving into a downstream task driven by foundational models. Learning robust and biologically interpretable cell representations from large-scale data represents a key direction for enhancing future model generalizability^41^.

## Methods

### Model architecture of HitAnno

HitAnno consists of three modules: a tokenization module, a representation module, and an annotation module. The tokenization module converts scATAC-seq data into cell sentences, a sequence-based representation constructed based on selected cell-type-specific peaks. The representation module employs a hierarchical transformer to parse the structured cell sentences at both peak level and peak-set level, learning cell embeddings. he annotation module maps the learned cell embeddings to cell type labels using an MLP. Given an scATAC-seq dataset, HitAnno formulates the annotation task based on a sparse cell-by-peak accessibility matrix, ***X*** ∈ ℝ^*n*×*p*^, where *n* and *p* denote the numbers of cells and peaks, respectively. In this matrix, the *i*-th row ***x***_*i*_ ∈ ℝ^*p*^ corresponds to the accessibility profile of cell *i*, with *x*_*ij*_ indicating the fragment count at peak *j*. Each cell *i* is associated with a cell-type label *y*_*i*_ ∈ {1, 2, ⋯, *c*}, where *c* denotes the number of cell types.

#### The tokenization module

For cell *i*, the tokenization module aims to convert scATAC-seq profile *x*_*i*_ into a sequence-based representation, termed cell sentence ***s***_*i*_. A cell sentence is composed of multiple clauses, each corresponding to a specific cell type in the reference dataset. Specifically, the clause is constructed based on a cell-type-specific peak set. Within a clause, tokens represent the binarized accessibility of peaks in the corresponding peak set. In addition, a special token [CLS_m_] is prepended to indicate the associated cell type. The *m*-th clause 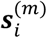 is represented as:

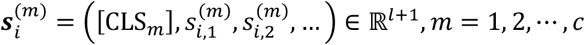

where 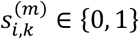 represents the binarized accessibility of the *k*-th peak in the *m*-th clause for cell *i*. The value of *l* controls the number of peaks included in each clause and thus determines the overall length of the cell sentence. It can be either manually specified or automatically determined based on the number of cell types *c* by default. In total, we obtain *c* clauses, which are concatenated in a fixed order to form the cell sentence ***s***_*i*_:

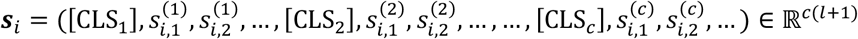

The construction of the cell sentence relies on cell-type-specific peak sets, which are identified for each cell type as follows. We first apply term frequency-inverse document frequency (TF-IDF) transformation to the raw count matrix ***X***. This transformation down-weights common peaks across cells^13, 42-44^. For each cell type, cells are divided into two subsets depending on whether they belong to that type, and a two-sided Welch’s t-test is performed for each peak on the TF-IDF-transformed matrix. Based on their resulting t-values, we rank the peaks in descending order. Peaks are then selected sequentially according to cell-type prevalence, starting from the rarest, with the top *l* previously unselected peaks chosen for each cell type. The peaks selected for each cell type form a corresponding peak set. Finally, we obtain *c* non-overlapping cell-type-specific peak sets. These peak sets are selected once during model training and fixed throughout subsequent inference.

#### The representation module

The objective of the representation module is to convert the structured cell sentence into a cell embedding. We generate two types of embeddings for the cell sentence: one capturing the chromatin accessibility profile, and another encoding the peak identity and positional context. First, the cell sentence ***s***_*i*_ is encoded via a dedicated embedding layer. Simultaneously, each token in the cell sentence is associated with a peak index, forming an index vector ***r***_*i*_ ∈ ℕ^*cl*^, which records the identity of each peak. This index vector is fed into a separate embedding layer. These two embeddings are then fused by element-wise addition, producing the cell sentence embedding ***E***_*i*_, which can be formulated as:

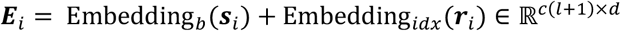

where Embedding_*b*_(·) and Embedding_*idx*_(·) denote the binary vector embedding layer and index vector embedding layer, respectively. In the embedding module, the dimensionality of the embeddings *d* is set to 128. As described in the tokenization module, each cell sentence ***s***_*i*_ is formed by *c* clauses 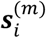. The result of 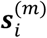 in the embedding layers can also be expressed as:

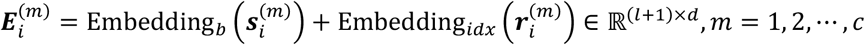

where 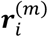 is the index vector corresponding to the specific peak set of *m*-th cell type for cell *i*.

In order to extract a representation that captures cell-type profiles, we employ the peak-level transformer and peak-set-level transformer to hierarchically parse the obtained cell sentence embedding to yield the cell embedding. The peak-level transformer is implemented using a single-layer BERT^45^ model, capturing local accessibility patterns among peaks in the same peak set. Each part 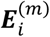 in the cell sentence is processed independently in parallel. In this level, the transformer performs self-attention within each cell-type-specific peak set, and the attention scores within each peak set are computed as follows:

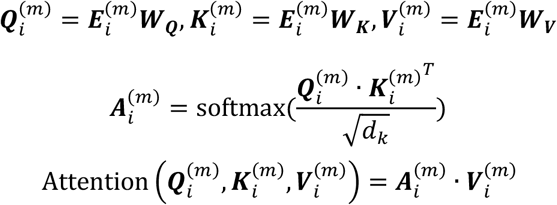

where, ***W***_***Q***_, ***W***_***K***_, and ***W***_***V***_ are learnable projection matrices for generating the query 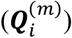, key 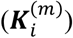, and value 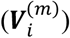 vectors, respectively; *d*_*k*._ denotes the dimensionality of the key vectors, and 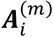 represents the resulting attention matrix. The relationship between the input 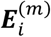 and the output 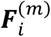 of the peak-level transformer is formulated as:

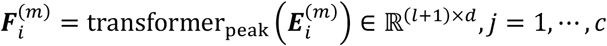

where transformer_peak_(·) denotes the peak-level transformer.

We extract the output vector corresponding to the [CLS_m_] token, 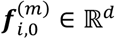, which serves as the overall embedding of the corresponding cell-type-specific peak set in cell *i*. Then we prepend a global [CLS] token embedding ***e***_*i*_ ∈ ℝ^*d*^ to the collection of peak-set embeddings. The resulting input to the peak-set-level transformer, denoted as ***G***_*i*_, is formulated as:

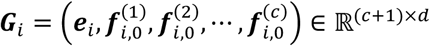

The peak-set-level transformer, also implemented as a single-layer BERT, models global dependencies across peak sets. The attention mechanism at this level operates over the entire collection of peak sets. The relationship between the input ***G***_*i*_ and output ***H***_*i*_ is formulated as:

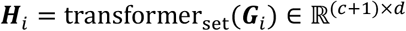

where transformer_set_(·) denotes the peak-set-level transformer.

We extract the output vector of the [CLS] token, ***h***_*i*,0_ ∈ ℝ^*d*^, as the cell embedding.

#### The annotation module

The annotation module is used to generate predicted cell type probabilities from the cell embedding. The cell embedding getting from the representation module is passed through a single-hidden-layer MLP for cell type annotation. The MLP has an input dimension of *d*, a hidden layer of size 64, and an output dimension equal to the number of cell types *c*. The resulting predicted label probability ***o***_*i*_ is formulated as:

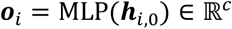

### Model training

We trained both the peak-level and peak-set-level transformer^46^ encoders using a single-layer BERT^45^ architecture with 128-dimensional embeddings and 16 attention heads. To accelerate attention computation, we employed FlashAttention-2^47^, and applied a learning rate scheduler with linear warmup for the first three epochs followed by linear decay. Training was conducted with the Adam^48^ optimizer (learning rate = 0.0001) and cross-entropy loss, using a batch size of 4 and mixed-precision computation with gradient scaling. The dataset was randomly split into training and validation sets at a 75:25 ratio, and early stopping was used based on validation accuracy, with training halted if no improvement was observed for three consecutive epochs (maximum 20 epochs). In practice, the models typically converged within 10 epochs, and the best-performing checkpoint was selected according to validation performance. In the atlas training task, a slightly larger set of cell-type specific peaks was used as HitAnno input; all other architectures and training settings remained unchanged.

### Baseline methods

We compared HitAnno with five baseline methods. Among them, PCA+SVM^10^ and MLP^16^ were originally developed for scRNA-seq-based cell type annotation, but were included due to their strong performance in recent benchmarking studies of transcriptomic classifiers^10, 16^. To adapt them for scATAC-seq data, we computed a gene activity score matrix by mapping peaks to genes based on their genomic proximity, using the “episcanpy.tl.geneactivity” function from the Episcanpy^49^ toolkit. For scATAC-seq-specific baselines, we selected EpiAnno^13^, Cellcano^14^, and SANGO^15^, which represent recent advances in chromatin accessibility-based annotation frameworks. One limitation of EpiAnno is that it does not handle batch effects and requires the entire dataset to be loaded into memory during probabilistic graph construction. To mitigate these limitations when working with large-scale datasets, we applied cell downsampling while preserving all other parameter settings. For all baseline methods, unless otherwise specified, model configurations and training procedures were consistent with those described in the original publications or followed the default settings provided by the authors. In ablation studies, TransformerMLP_acc and TransformerMLP replace the hierarchical two-level transformer with a single transformer layer. Instead of the cell-type-specific peak selection strategy, HitAnno_acc and TransformerMLP_acc select globally the most accessible peaks based on accessibility rankings.

### Model evaluation

To evaluate the annotation performance of our model, we adopted three metrics: accuracy, macro-F1 score, and Cohen’s kappa score. Accuracy directly measures the overall correctness of the model across all cell types. However, in imbalanced annotation scenarios, the accuracy may fail to reflect performance on rare cell types. To address this, we included the macro-F1 score, which calculates the unweighted average F1 score across all classes, thus highlighting the ability of model to annotate both major and rare cell types. Additionally, we employed Cohen’s kappa score to account for agreement occurring by chance. This metric provides a more robust measure of performance under class imbalance by quantifying the consistency between predicted and true labels beyond random chance. The detailed formulas for computing these metrics are available in Supplementary Note 1. To test the effect of cell type imbalance, we individually sampled cells from different categories to simulate various imbalance degrees, following prior works^20, 42^. When targeting major cell types, we downsampled those whose proportions exceeded 5%; otherwise, we downsampled the remaining rare cell types. For both downsampling and dropout experiments, we adopted a progressive sampling strategy. This ensured consistent comparability across different levels of controlled data modification. For all ablation experiments, models were run on the same GPU device with 24 GB of memory to control for hardware variability.

### Model interpretation analysis

To further investigate the interpretability of our model, we extracted attention matrices from the trained single-layer transformer encoders. For each annotated cell, we collected the attention weights from this layer and averaged them across all heads to obtain a unified representation of peak-to-peak (or peak-set-to-peak-set) attention. When multiple cells were involved, we further averaged the attention matrices across samples to derive the overall attention pattern. To infer cis-regulatory chromatin interactions, we used Cicero^23^ to predict cis-regulatory chromatin interactions for each cell type. We preprocessed scATAC-seq data for each cell type individually, using default parameters of the “detect_genes”, “estimate_size_factors”, “preprocess_cds”, and “reduce_dimension” functions, followed by conversion into Cicero-compatible objects via “make_atac_cds”. Chromatin interactions were then inferred, using the “run_cicero” function.

### Dataset collection and preprocessing

The scATAC-seq datasets used in this study were obtained from their original sources (see Data availability). Peak coordinates were converted to the hg38 reference genome using UCSC LiftOver^50^. For each dataset, we directly adopted the cell-by-peak accessibility matrix provided by the original studies, without additional peak calling or read alignment. Only basic filtering of low-quality cells, as defined in the original publications, was applied.

## Data availability

All datasets used in this study are publicly available and were obtained from previously published studies. The Domcke2020^17^ dataset is available from the Gene Expression Omnibus (GEO) under accession number GSE149683. The Herring2022^25^ dataset can be accessed via GEO accession GSE168408, and the Li2023^29^ dataset is available under GEO accession GSE244618 as well as from http://catlas.org. The Trevino2021^24^ dataset is accessible through GEO accession GSE162170, and the Zhang2021^18^ dataset is available under GEO accession GSE184462. The Ziffra2021^26^ dataset can be accessed at https://cortex-atac.cells.ucsc.edu. The Kanemaru2023^30^ dataset can be accessed at https://www.heartcellatlas.org/. The Li2023b^31^ dataset is available under GEO accession GSE219281. The human adrenal tissue Hi-C data used for model interpretation analysis are publicly available from the ENCODE project^51^ under accession number ENCFF499BVX.

## Code availability

The source code of HitAnno is available on Github (https://github.com/zianwang-bioinfo/HitAnno/).

## Acknowledgements

This work was partly supported by the National Key Research and Development Program of China (grant nos. 2025YFC3409300, 2023YFF1204802), the National Natural Science Foundation of China (grant nos. 32550616, 62273194) and Beijing Natural Science Foundation (grant no. L242026).

## Author contributions

R.J. conceptualized the study and provided overall supervision. Z.W. and X.C. jointly developed and evaluated the HitAnno. Z.W., X.C., and K.L. performed data collection and preprocessing. X.C., Z.G., and Z.L. contributed to the refinement of HitAnno. The manuscript was drafted by Z.W., X.C., and R.J., with feedback and revisions from all authors.

