## Supplementary Information for "HitAnno: Atlas-level cell type annotation based on scATAC-seq data via a hierarchical language model"

### Content


**Supplementary Notes** ______ 3

**Supplementary Note 1: Formulas for all the metrics** ______ 3

**Supplementary Figures** ______ 5

**Supplementary Figure 1** ______ 5

**Supplementary Figure 2** ______ 6

**Supplementary Figure 3** ______ 7

**Supplementary Figure 4** ______ 8

**Supplementary Figure 5** ______ 9

**Supplementary Figure 6** ______ 10

**Supplementary Figure 7** ______ 11

**Supplementary Figure 8** ______ 12

**Supplementary Figure 9** ______ 13

**Supplementary Figure 10** ______ 14

**Supplementary Figure 11** ______ 15

**Supplementary Tables** ______ 16

**Supplementary Table 1** ______ 16

### Supplementary Notes

#### Supplementary Note 1: Formulas for all the metrics

This section describes the formulas used to compute the evaluation metrics reported in this study. Let $n$ denote the number of cells. For each sample $i$ $(i = 1, 2, \ldots, n)$, let $y_i$ denote the ground truth label and $\hat{y}_i$ denote the predicted label. Let $n_{ct}$ denote the number of cell types in the dataset.

Accuracy is calculated as the proportion of correct predictions among all samples:

$$\text{accuracy} = \frac{\sum_{i=1}^{n} I(y_i = \hat{y}_i)}{n}$$

where $I(\text{condition})$ is the indicator function, equal to 1 if the condition is true and 0 otherwise.

Macro-F1 score (macro-f1) is the unweighted average of per-class F1 scores. For each class $c$, precision ($P_c$) and recall ($R_c$) are defined as:

$$P_c = \frac{\text{TP}_c}{\text{TP}_c + \text{FP}_c}$$

$$R_c = \frac{\text{TP}_c}{\text{TP}_c + \text{FN}_c}$$

where $\text{TP}_c$, $\text{FP}_c$, and $\text{FN}_c$ denote the total true positives, false positives, and false negatives of class $c$. And the F1 score for class $c$ is then:

$$\text{F1}_c = \frac{2P_cR_c}{P_c + R_c}$$

The macro-f1 is given by:

$$\text{macro f1} = \frac{1}{n_{ct}} \sum_{c=1}^{n_{ct}} \text{F1}_c$$

Cohen's kappa score ($\kappa$) measures the agreement between predicted and true labels, adjusted for chance. The observed agreement ($p_o$) is defined as:

$$p_o = \text{accuracy}$$

The expected agreement ($p_e$) is computed as:

$$p_e = \sum_{c=1}^{n_{ct}} p_y^{(c)} p_{\hat{y}}^{(c)}$$

where $p_y^{(c)}$ and $p_{\hat{y}}^{(c)}$ denote the proportions of class c in the true and predicted labels, respectively, summed over all classes $c = 1, 2, \ldots, n_{ct}$. The $\kappa$ is given by:

$$\kappa = \frac{p_o - p_e}{1 - p_e}$$

Imbalance degree quantifies the deviation of the label distribution from a perfectly uniform distribution across cell types. The label entropy is computed as:

$$H = -\sum_{c=1}^{n_{ct}} p_y^{(c)} \log(p_y^{(c)})$$

The imbalance degree is then defined as:

$$\text{Imbalance degree} = 1 - \frac{H}{\log(n_{ct})}$$

This metric ranges from 0 (perfectly balanced) to 1 (maximally imbalanced, where all samples belong to a single class).

### Supplementary Figures

#### Supplementary Figure 1

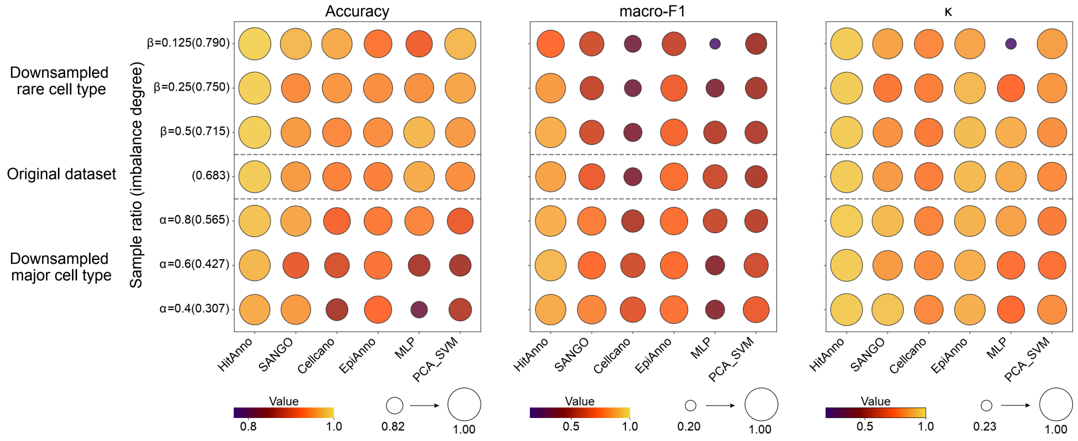


**Supplementary Figure 1.** Accuracy, macro-F1, and $\kappa$ of HitAnno and five baseline methods under varying levels of cell type imbalance.

#### Supplementary Figure 2

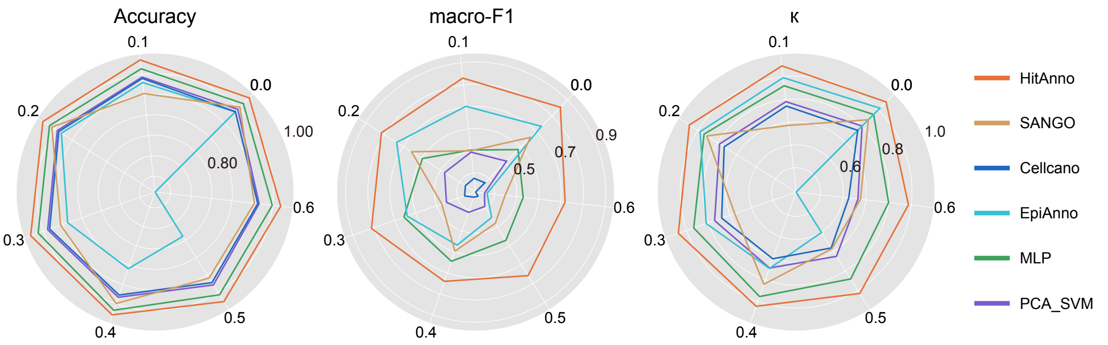


**Supplementary Figure 2.** Accuracy, macro-F1, and $\kappa$ of HitAnno and five baseline methods under varying levels of dropout rates.

#### Supplementary Figure 3

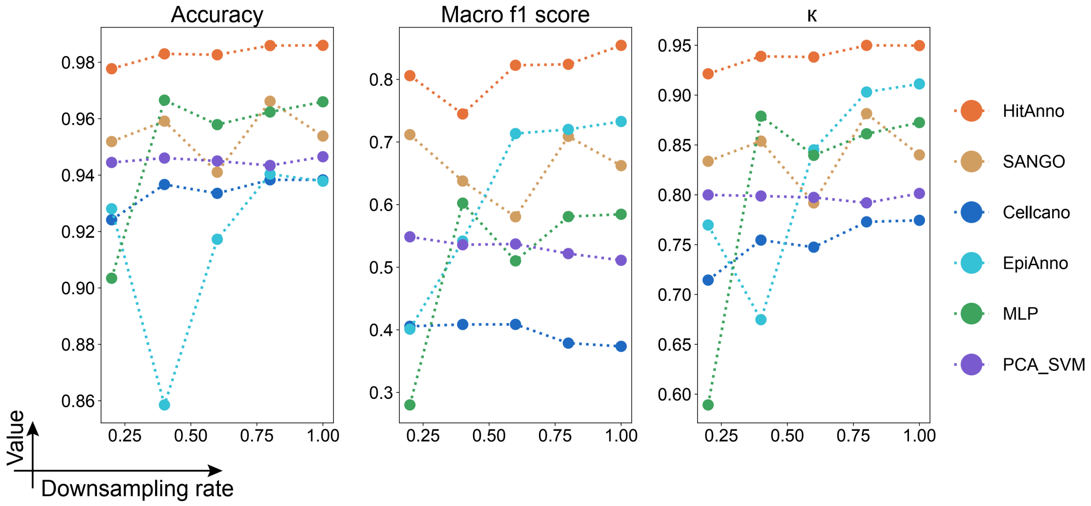


**Supplementary Figure 3.** Accuracy, macro-F1, and $\kappa$ of HitAnno and five baseline methods under varying levels of downsampling ratios.

#### Supplementary Figure 4

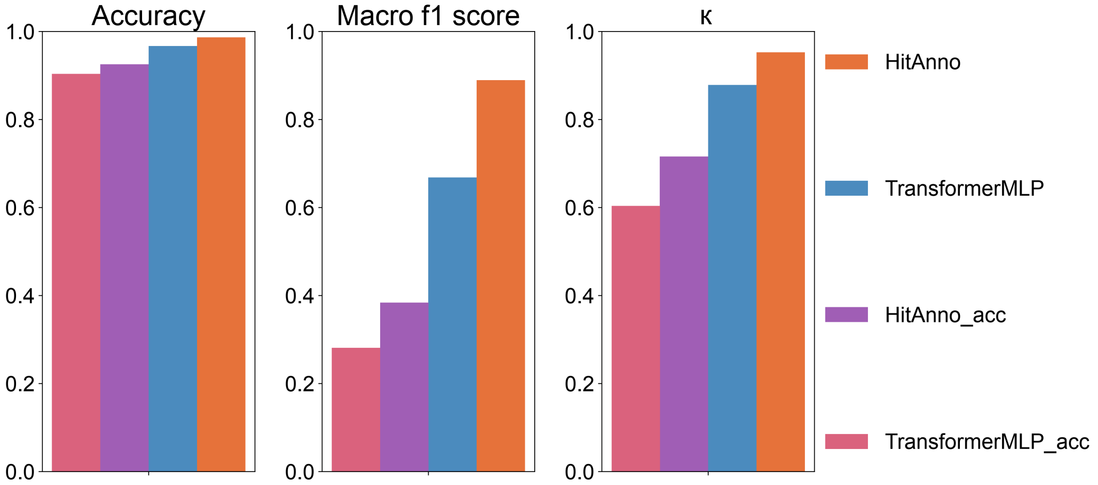


**Supplementary Figure 4.** Bar plots showing the Accuracy, macro-F1, and $\kappa$ of HitAnno after removing individual modules.

### Supplementary Figure 5

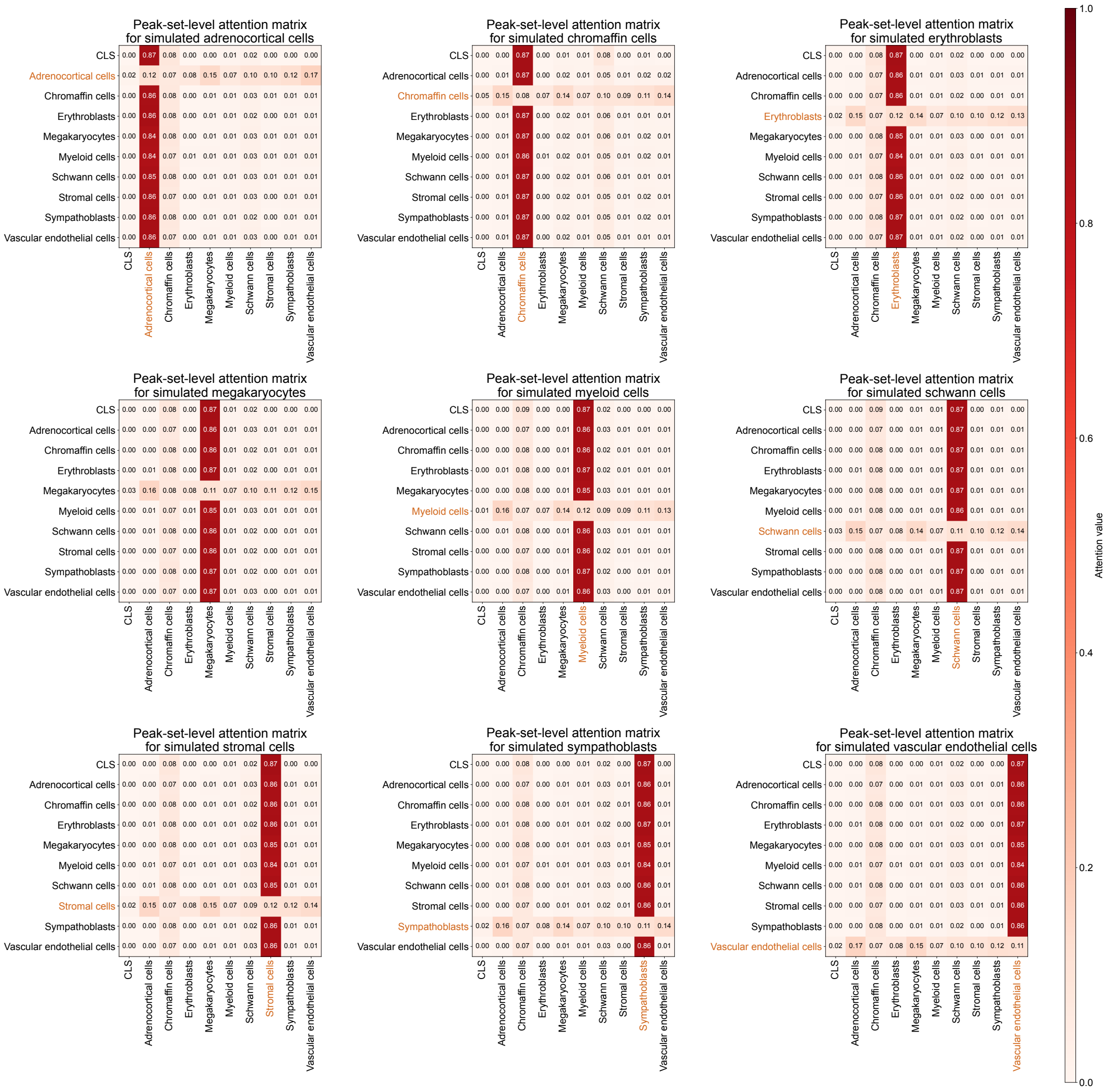


**Supplementary Figure 5.** Attention matrix at the peak-set level from the trained model. The input consists of artificially constructed binary vectors, in which only the cell-type-specific peaks of one cell type are set to 1, and all other peaks are set to 0.

#### Supplementary Figure 6

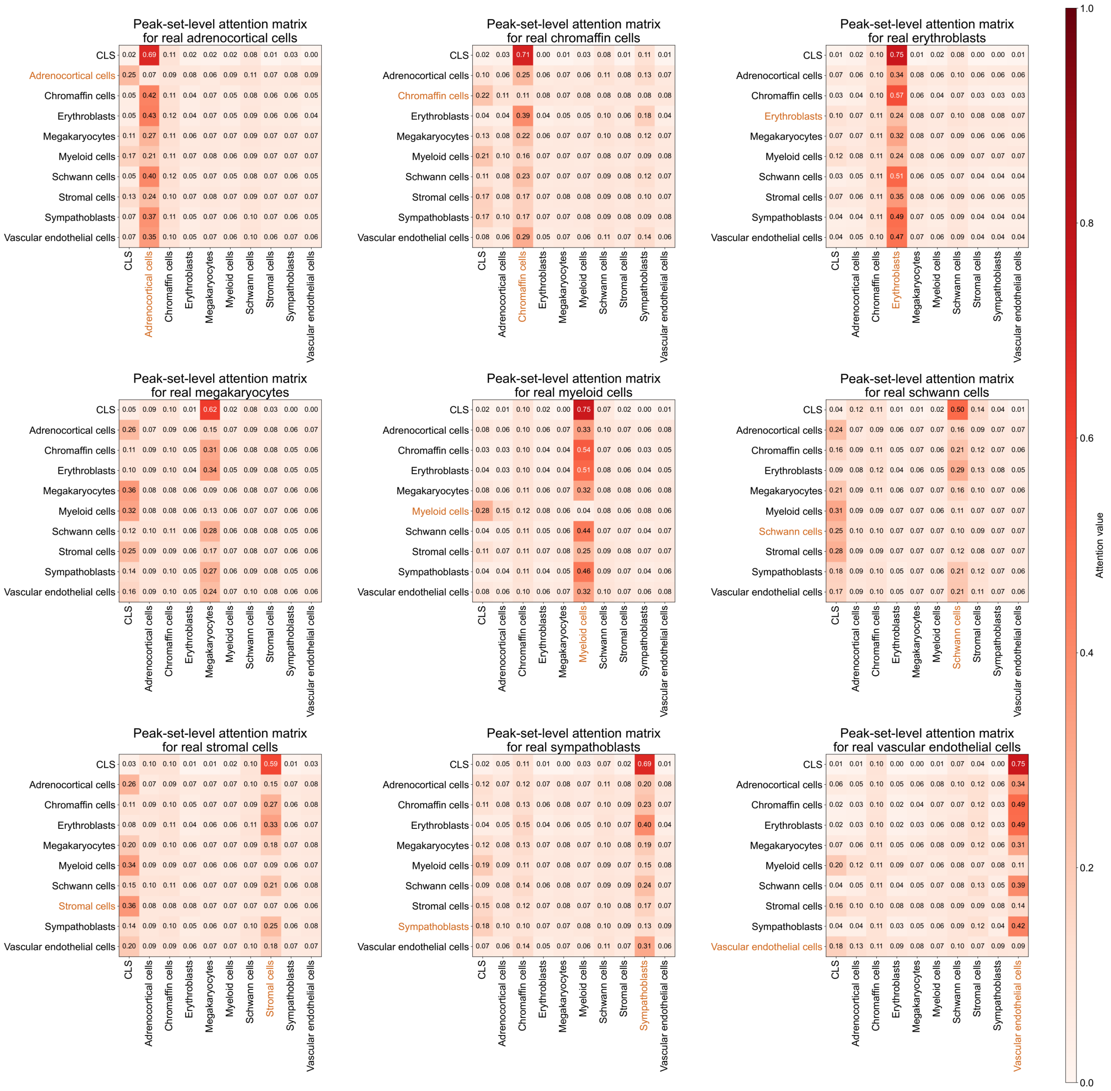

**Supplementary Figure 6.** Attention matrix at the peak-set level from the trained model. Each subpanel shows the attention output for a specific cell type in the dataset, averaged first across attention heads and then across all cells of that type.

#### Supplementary Figure 7

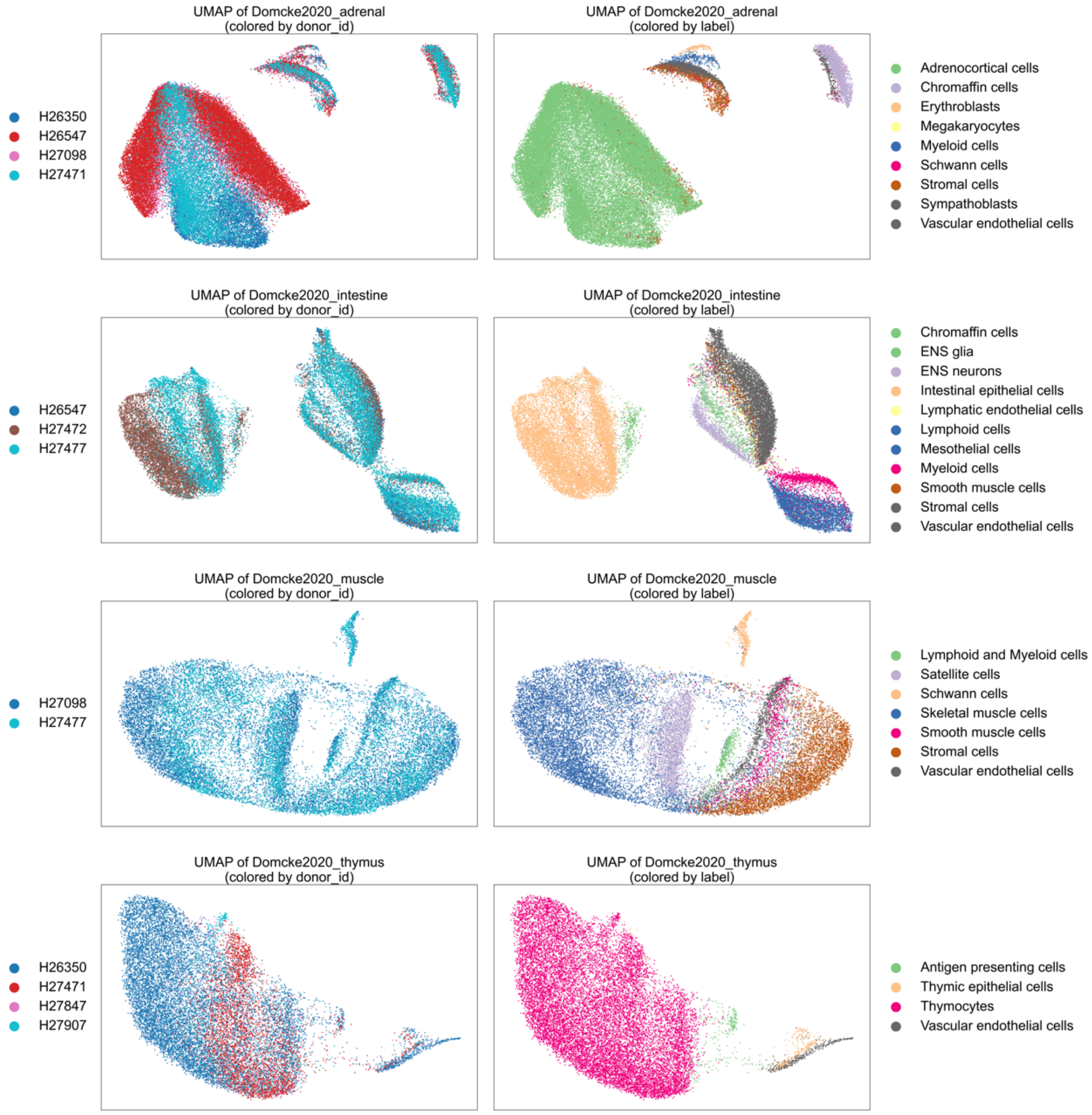


**Supplementary Figure 7.** UMAP visualizations of four datasets (Domcke2020_adrenal, Domcke2020_intestine, Domcke2020_muscle, and Domcke2020_thymus). For each dataset, two panels are shown: one colored by donor ID and the other by cell type label.

#### Supplementary Figure 8

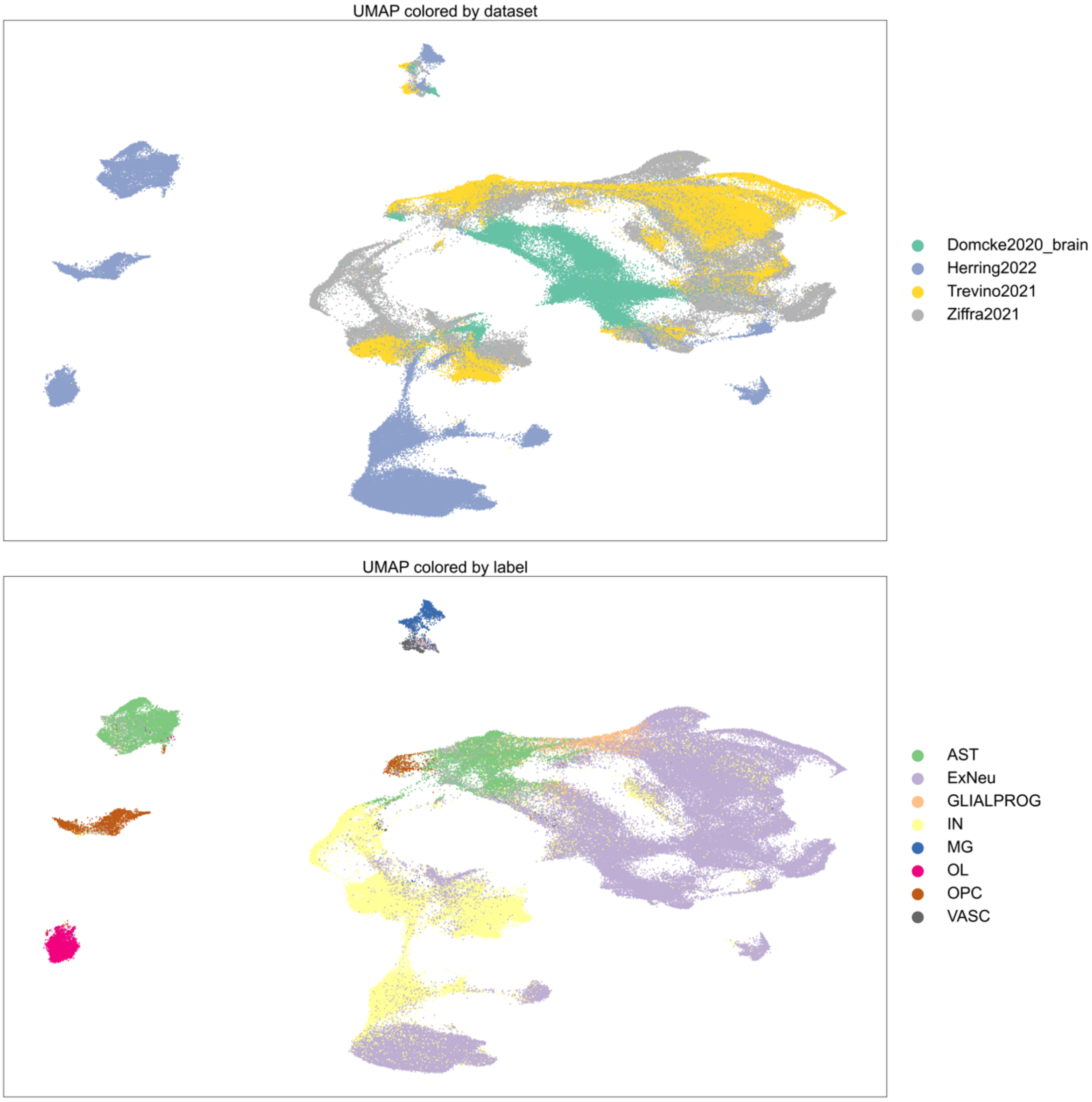


**Supplementary Figure 8.** UMAP visualizations of an integrated dataset comprising four studies (Domcke2020_brain, Herring2022, Trevino2021, and Ziffra2021). Two panels are shown: one colored by dataset origin and the other by cell type label.

#### Supplementary Figure 9

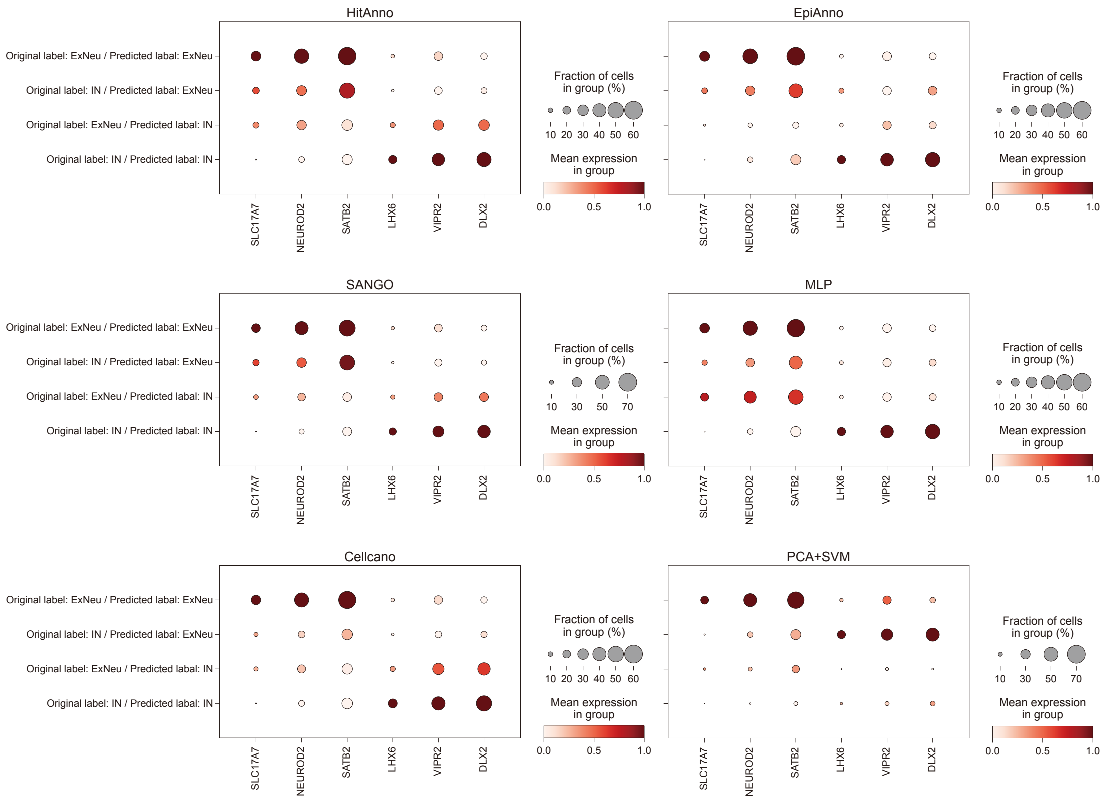


**Supplementary Figure 9.** Gene activity scores of marker genes for ExNeu and IN in four categories of cells based on original and predicted labels. Dot color represents the average gene activity score, and dot size reflects the fraction of cells in each category expressing the corresponding marker gene.

**Supplementary Figure 10**

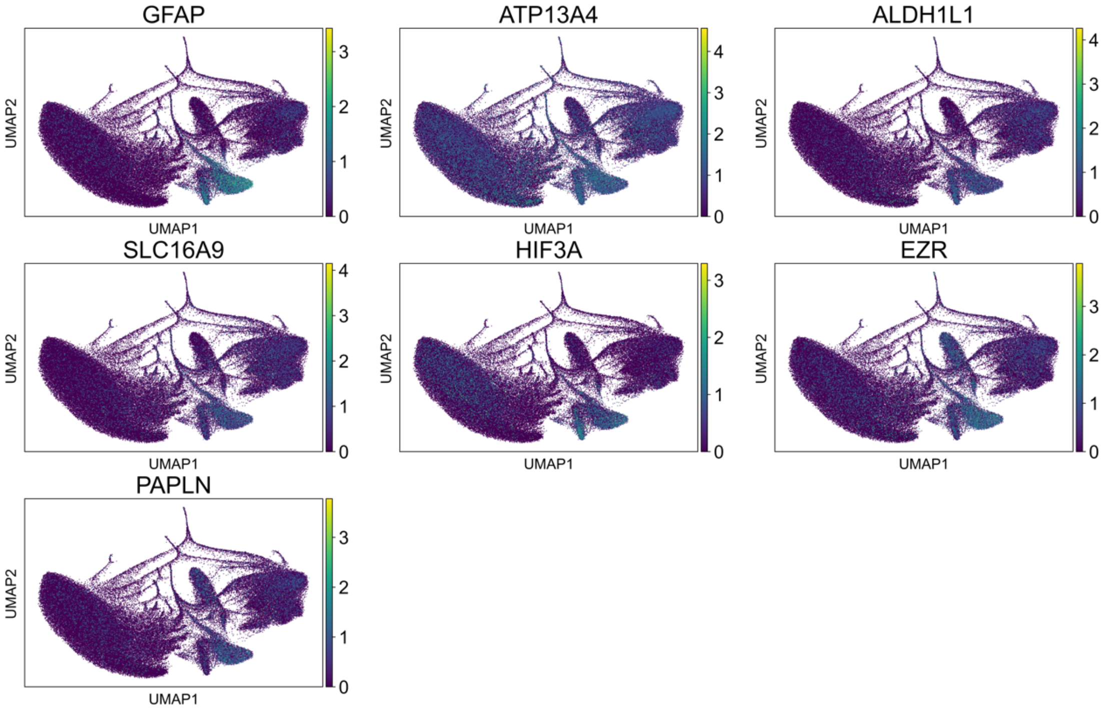


**Supplementary Figure 10.** UMAP visualizations of the Li2023b dataset. The panels show the gene activity scores of the astrocyte marker genes.

#### Supplementary Figure 11

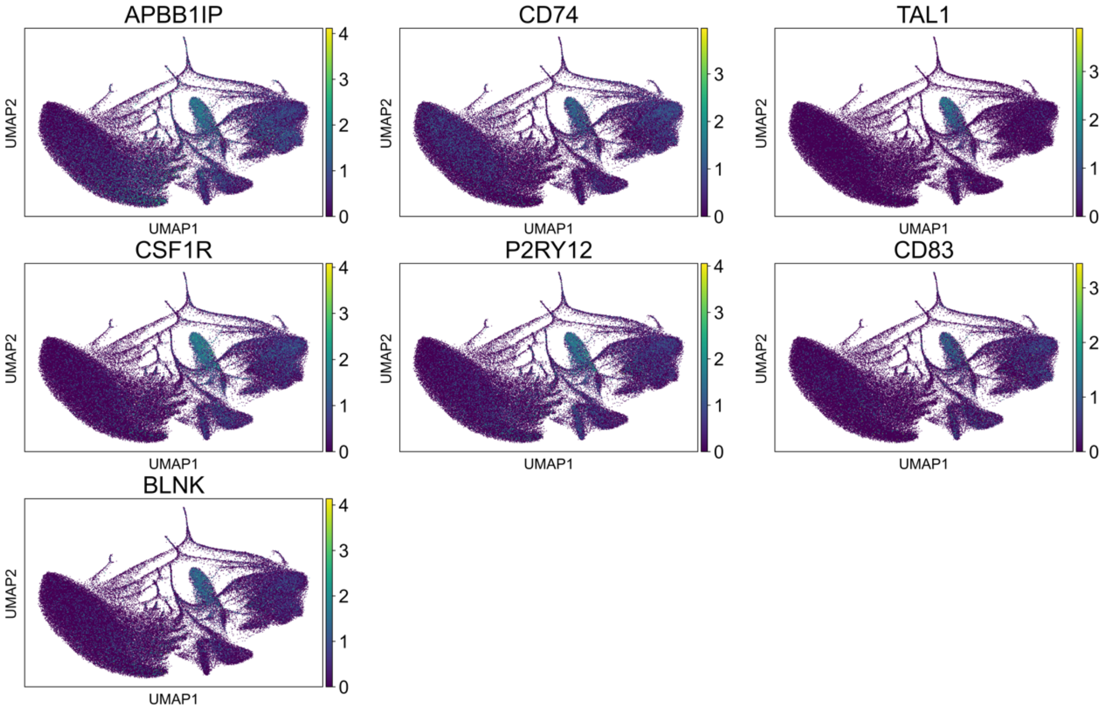


**Supplementary Figure 11.** UMAP visualizations of the Li2023b dataset. The panels show the gene activity scores of the microglia marker genes.

#### Supplementary Tables

##### Supplementary Table 1

| Dataset | Number of cells | Number of peaks | Number of cell types | Sequencing depth | Sparsity (%) | Tissue | Sequencing technology |
|---|---|---|---|---|---|---|---|
| Herring2022 | 85846 | 190134 | 8 | 3865.9 | 98.176 | Brain tissue | 10X |
| Trevino2021 | 39268 | 190134 | 7 | 18431.6 | 95.709 | Brain tissue | 10X |
| Ziffra2021 | 75256 | 190134 | 7 | 5662.2 | 97.050 | Brain tissue | 10X |
| Domcke2020_adrenal | 63958 | 1050819 | 9 | 7599.8 | 99.601 | Adrenal gland | sci-ATAC-seq3 |
| Domcke2020_brain | 84520 | 190134 | 8 | 2384.6 | 99.354 | Brain tissue | sci-ATAC-seq3 |
| Domcke2020_intestine | 42942 | 1050819 | 13 | 7553.5 | 99.619 | Intestinal tissue | sci-ATAC-seq3 |
| Domcke2020_muscle | 27181 | 1050819 | 10 | 10541.9 | 99.447 | Muscle tissue | sci-ATAC-seq3 |
| Domcke2020_thymus | 21499 | 1050819 | 4 | 9173.6 | 99.574 | Thymic tissue | sci-ATAC-seq3 |
| Zhang2021_artery | 46576 | 1154464 | 19 | 11885.9 | 99.294 | Vascular tissue | sci-ATAC-seq |
| Zhang2021_liver | 6328 | 1154464 | 6 | 8344.7 | 99.536 | Hepatic tissue | sci-ATAC-seq |
| Zhang2021_lung | 41188 | 1154464 | 22 | 6867.8 | 99.601 | Pulmonary tissue | sci-ATAC-seq |
| Zhang2021_pancreas | 32887 | 1154464 | 20 | 6769.1 | 99.612 | Pancreatic tissue | sci-ATAC-seq |
